# Genome-wide screen for enhanced noncanonical amino acid incorporation in yeast

**DOI:** 10.1101/2022.04.28.489958

**Authors:** Matthew T. Zackin, Jessica T. Stieglitz, James A. Van Deventer

**Affiliations:** Chemical and Biological Engineering Department, Tufts University, Medford, Massachusetts 02155, USA; Biomedical Engineering Department, Tufts University, Medford, Massachusetts 02155, USA

**Keywords:** Noncanonical amino acids, yeast knockout collection, fluorescence-activated cell sorting, aminoacyl-tRNA synthetases, amber suppression

## Abstract

Numerous applications of noncanonical amino acids (ncAAs) in basic biology and therapeutic development require efficient protein biosynthesis using an expanded genetic code. However, achieving such incorporation at repurposed stop codons in cells is generally inefficient and limited by complex cellular processes that preserve the fidelity of protein synthesis. A more comprehensive understanding of the processes that contribute to ncAA incorporation would aid in the development of genomic engineering strategies for augmenting genetic code manipulation. In this work, we screened a pooled *Saccharomyces cerevisiae* molecular barcoded yeast knockout (YKO) collection to identify single-gene knockout strains exhibiting improved ncAA incorporation efficiency in response to the amber (TAG) stop codon. We used a series of intracellular fluorescent reporters in tandem with fluorescence activated cell sorting (FACS) to identify 55 unique candidate deletion strains. Identified genes encode for proteins that participate in diverse cellular processes; many of the genes have no known connection with protein translation. We then verified that two knockouts, *yil014c-aΔ* and *alo1Δ*, had improved incorporation efficiency using independently acquired strains possessing the knockouts. Characterizations of the activity of *yil014c-aΔ* and *alo1Δ* with additional orthogonal translation systems and ncAAs indicate that deletion of each of these genes enhances ncAA incorporation efficiency without loss of fidelity over a wide range of conditions. Our findings highlight opportunities for further modulating gene expression with genetic, genomic, and synthetic biology approaches to improve ncAA incorporation efficiency. In addition, these discoveries have the potential to enhance our fundamental understanding of protein translation. Ultimately, this study provides a foundation for future efforts to engineer cells to incorporate ncAA at greater efficiencies, which in turn will streamline the realization of applications utilizing expanded genetic codes ranging from basic biology to drug discovery.

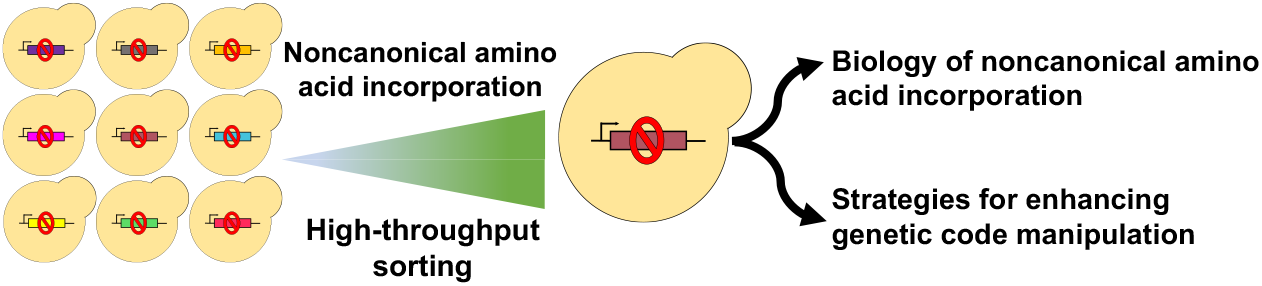

## Introduction

Genetic encoding of noncanonical amino acids (ncAAs) enables new approaches for better understanding, controlling, and engineering protein properties (*1*). NcAAs possess diverse properties, including biorthogonal groups (*2, 3*) and fluorescent probes (*4, 5*), that can be exploited for a wide range of applications such as furthering understanding of post-translational modifications (*6–8*) and biological therapeutic development (*9–11*). Effective employment of ncAAs for these applications is contingent on their efficient incorporation at repurposed stop codons or other “blank” codons not normally encoding for amino acids. Stop codon suppression for ncAA incorporation is limited by complex cellular processes and machinery that preserve the fidelity of protein synthesis. Efforts to improve ncAA incorporation by engineering the cellular machinery directly implicated in protein translation, including ribosomes and release factors, have led to enhanced genetic code manipulation in *E. coli* (*12–14*). Engineering orthogonal translation systems (OTSs), comprised of a suppressor tRNA and an aminoacyl-tRNA synthetase (aaRS), has further augmented genetic code expansion by increasing the number of different ncAAs encodable into proteins and improving incorporation efficiencies (*15–18*). Ongoing efforts to compress the *E. coli* genome to 61 or fewer codons demonstrate that genome engineering is a viable strategy for creating cells that support efficient protein biosynthesis with expanded genetic codes (*19, 20*).

In addition to complete genome reengineering, the tools of genetics, genomics, and synthetic biology provide opportunities to utilize more targeted strategies to enhance the production of proteins containing ncAAs. Manipulation of gene expression via overexpression, knockdowns, and knockouts are common approaches to improve the yield of biosynthesized compounds (*21*). Applying this strategy to the production of proteins containing ncAAs poses challenges due to the essentiality of the components of the protein translation apparatus. Beyond engineering the genes directly related to protein translation, there are numerous opportunities to modulate the expression and function of genes that indirectly impact ncAA incorporation. However, the identities of these genes, let alone the roles of their protein products in ncAA incorporation, are not fully known or understood. A more comprehensive understanding of the processes that contribute to ncAA incorporation would aid in the development of genomic engineering strategies for augmenting genetic code manipulation. To our knowledge, there has never been a genome-wide screen to identify processes involved in ncAA incorporation. However, the tools for such a study are available: pooled yeast knockout (YKO) collections, comprised of ~6,000 knockout strains that cover approximately 96% of the yeast genome, have enabled genome-wide studies to understand genes implicated in various aspects of eukaryotic biology (*22, 23*). YKO collections have also facilitated genome-wide functional profiling in various growth stress conditions and studies for identifying genes and protein involved within cellular pathways, such as genes that regulate the yeast mating pathway (*24–26*). In addition, there exist multiple ways to analyze ncAA incorporation quantitatively and qualitatively in yeast using intracellular fluorescent protein reporters in high throughput (*27–29*).

In this work, we screened a pooled *Saccharomyces cerevisiae* heterozygous diploid BY4743 molecular barcoded yeast knockout (YKO) collection to identify single-gene knockout strains exhibiting improved ncAA incorporation efficiency in response to the amber (TAG) stop codon (*22*). We used a series of intracellular fluorescent reporters in tandem with fluorescence activated cell sorting (FACS) to identify 55 unique candidate deletion strains. Identified genes encode for proteins that participate in diverse cellular processes; many of the genes have no known connection with protein translation, with several genes having uncharacterized functions. We then verified that two knockouts, *yil014c-aΔ* and *alo1Δ*, had improved incorporation efficiency using independently acquired strains possessing the knockouts. Finally, we characterized the activity of *yil014c-aΔ* and *alo1Δ* with additional orthogonal translation systems and ncAAs and found that deletion of each of these genes enhances ncAA incorporation efficiency without loss of fidelity for a number of conditions. Our findings highlight opportunities for further modulating gene expression with genetic, genomic, and synthetic biology approaches to improve ncAA incorporation efficiency. In addition, these discoveries have the potential to enhance our fundamental understanding of protein translation in eukaryotic systems. Ultimately, this study provides a foundation for future efforts to engineer cells to incorporate ncAA at greater efficiencies, which in turn will streamline the realization of applications utilizing expanded genetic codes ranging from basic biology to drug discovery.

## Results and Discussion

### Screening a pooled yeast knockout collection

To identify gene deletions in yeast that enhance the efficiency of noncanonical amino acid (ncAA) incorporation at repurposed stop codons, a genome-wide screen was performed with the pooled *S. cerevisiae* heterozygous diploid BY4743 molecular barcoded yeast knockout (YKO) collection. The YKO collection consists of ~6,000 unique single-gene deletion strains that contain unique DNA barcodes that enable the identification of the specific gene deletion present in each strain of the collection (*26, 30*). Prior to screening, a fluorescent protein reporter and an orthogonal translation system (OTS) were transformed via electroporation into the YKO collection. Four combinations of reporters and OTSs enabled diverse screening conditions to increase the chance of discovering gene knockouts with true enhancement of ncAA incorporation (SI Table 1). Each unique transformed library was comprised of one of two reporters, BXG or BXG-altTAG, in combination with one of two OTSs, LeuOmeRS/tRNA_CUA_^Leu^ or TyrOmeRS/tRNA_CUA_^Tyr^ (Figure 1A). Both reporters encode blue fluorescent protein (BFP) linked to green fluorescent protein (GFP) with an encoded peptide linker containing an amber stop codon. The two reporters have different stop codon positions that have previously been shown to exhibit different readthrough efficiencies (*27*). The reporter architecture enables high-throughput evaluations of ncAA incorporation efficiency at the level of both individual clones and whole populations; the reporters are also compatible with fluorescence-activated cell sorting (FACS) (*29*). The OTS is the machinery that enables ncAA incorporation at amber stop codons and consist of an aminoacyl-tRNA synthetase (aaRS) and a suppressor tRNA_CUA_. Both of the OTSs utilized in screening support the insertion of OmeY in response to the amber stop codon (*28*). All transformed YKO libraries were evaluated for transformation efficiency and strain diversity prior to screening (SI Table 2). All library transformations yielded at least 3.0 × 10^4^ transformants, which is 5-fold greater than the 6000-member YKO collection. Strain diversity in transformed collections was evaluated by isolating 8–10 colonies per library and identifying their associated gene deletion by performing Sanger sequencing of the molecular barcodes. In all four libraries, no more than one deletion strain was repeated (SI Table 3), indicating that the libraries possessed sufficient diversity to support identification of strains of interest via high-throughput screening.

**Figure 1.**
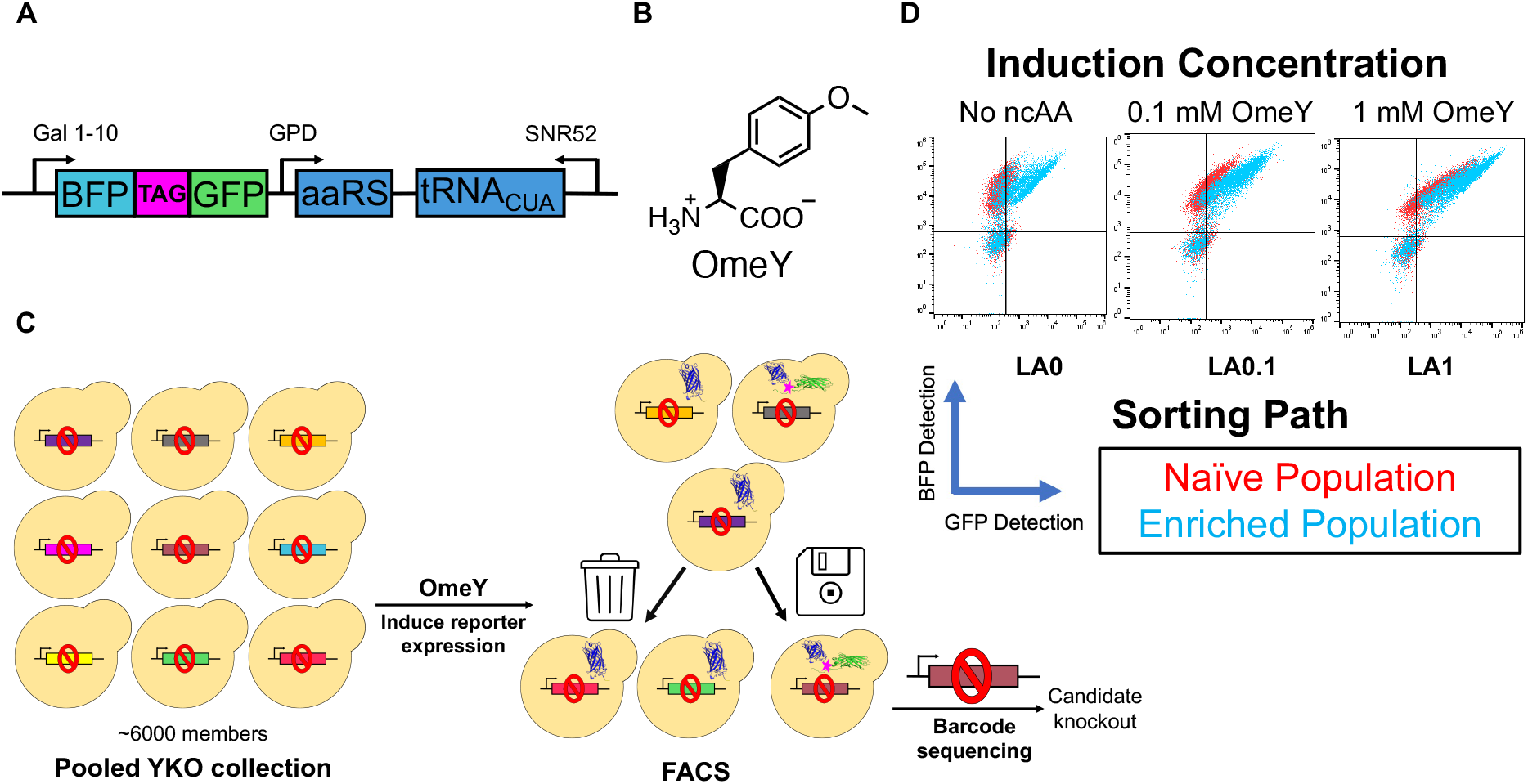
Overview of screening campaign. (A) Diagram of the general architecture of the four unique plasmid systems transformed into the ~6,000-member yeast knockout (YKO) collection containing the reporter and orthogonal translation system (OTS). The reporters consisted of a BFP linked to GFP by a peptide linker containing an amber (TAG) stop codon. Two reporters, BXG and BXG-altTAG, that differ by the location of the stop codon within the peptide linker were used. The OTS contained an aaRS and suppressor tRNA_CUA_. Two aaRSs, TyrOmeRS and LeuOmeRS, were used. (B) Structure of ncAA, *O*-methyl-L-tyrosine (OmeY), used for screens. (C) General schematic of the screening process. A pooled YKO collection transformed with the reporter/OTS was induced in the presence of 0, 0.1, or 1 mM OmeY. FACS was used to isolate cells with high GFP expression, which indicated successful OmeY incorporation in the reporter (or, in some cases, canonical amino acid incorporation). Following multiple rounds of screening, the barcodes of isolated colonies from enriched populations were Sanger sequenced to identify their respective gene deletions. (D) Representative flow cytometry overlay dot plots comparing readthrough of naïve (presort) and enriched (post sort) populations of three sorting paths. Populations were induced with the same concentration of OmeY used for screening.

The YKO libraries were screened via FACS following induction in the presence of the ncAA *O*-methyl-L-tyrosine (OmeY) (Figure 1B), or in the absence of ncAA. Libraries were screened under multiple conditions—different OmeY induction concentrations, different fluorescent detection approaches, and number of rounds of screening—in order to identify a wide range of candidate deletion strains with potentially enhanced ncAA incorporation efficiency (SI Table 4). In total, screens were performed utilizing 12 separate tracks (SI Table 4). Our screening strategy employed only positive sorts, where acquisition gates were drawn to collect the cells with highest C-terminal (i.e., GFP) reporter detection (Figure 1C). Following FACS, the ncAA incorporation efficiency of the enriched and the naïve populations were qualitatively analyzed and compared via analytical flow cytometry. After two or three rounds of sorting, the enriched populations exhibited an observable increase in GFP detection compared to respective naïve populations based on the flow cytometry plots (Figure 1D). This indicates that our screening strategy was capable of isolating enriched populations of deletion strains with enhanced GFP fluorescence, suggesting potential increases in ncAA incorporation efficiency in those cells. Interestingly, even following induction in the absence of ncAAs, the enriched populations exhibited higher levels of stop codon readthrough. These trends of improved ncAA incorporation (or stop codon readthrough in the absence of ncAAs) were observed across all screens conducted here (SI Figure 1).

### Identification and initial characterization of candidate deletion strains

After screening the YKO libraries and qualitatively evaluating the enriched populations, we sought to identify and characterize the individual deletion strains. From the 12 screening paths, we performed Sanger sequencing of the molecular barcodes of 104 isolated knockout strains, which yielded 55 unique deletions (SI Table 5). Genes of particular interest, based on either their presence in multiple sorting paths or their dominance in a specific sorting path, include *YIL014C-A* (also known as *YIL015C-A*), *ALO1, YKL131W, ABZ2, IRC24, MGA1, GPB1, RFA3*, and *APL1* (Table 1). According to the *Saccharomyces* Genome Database (SGD), the identified deletions are involved in a wide range of cellular processes, many of which are not currently known to be involved in processes directly relevant to protein biosynthesis (*31*). From an SGD gene ontology analysis, commonly identified cellular process groups included transcription (nine genes), metabolism (eight genes), cell cycle (eight genes), response to chemical stimuli (seven genes), and transport (seven genes) (see *Materials and Methods* for explanation on groupings). Fourteen genes, including the most frequently identified gene deletion in our sorts (*YIL014C-A*), encode for proteins with unknown or uncharacterized functions (Table 2).

**Table 1.** List of frequently identified gene deletions including which screens the gene deletion strain was identified in and the *Saccharomyces* Genome Database (SGD) description of the genes of interest (taken directly from the database) (*31*). Genes listed here were either identified in multiple sorting paths or were the dominant deletion identified in one sorting path.

| Systematic Name | Standard Name | Screen(s) Identified | Description (from the SGD) |
| --- | --- | --- | --- |
| YIL014C-A | N/A | LA0, LA0.1, LA1, 488_0, YA1 | Putative protein of unknown function |
| YML086C | *ALO1* | LA0.1, LA1 | D-Arabinono-1,4-lactone oxidase; catalyzes the final step in biosynthesis of dehydro-D-arabinono-1,4-lactone, which is protective against oxidative stress |
| YKL131W | N/A | LA0.1, 488_0 | Dubious open reading frame; unlikely to encode a functional protein |
| YMR289W | *ABZ2* | 488_0.1 | Aminodeoxychorismate lyase (4-amino-4-deoxychorismate lyase); catalyzes the third step in para-aminobenzoic acid biosynthesis; involved in folic acid biosynthesis |
| YIR036C | *IRC24* | LX0.1 | Putative benzyl reductase |
| YGR249W | *MGA1* | 488_0, YX0.1 | Protein similar to heat shock transcription factor |
| YKL131W | N/A | LA0.1, 488_0 | Dubious open reading frame; unlikely to encode a functional protein |
| YOR371C | *GPB1* | 488_1, YA0.1 | Multistep regulator of cAMP-PKA signaling; inhibits PKA downstream of Gpa2p and Cyr1p, thereby increasing cAMP dependency; promotes ubiquitin-dependent proteolysis of Ira2p |
| YJL173C | *RFA3* | LA0 | Subunit of heterotrimeric Replication Protein A (RPA); RPA is a highly conserved single-stranded DNA binding protein complex involved in DNA replication, repair, recombination; |
| YJR005W | *APL1* | 488_1, YA0.1 | Beta-adaptin; large subunit of the clathrin associated protein complex (AP-2); involved in vesicle mediated transport |

**Table 2.**
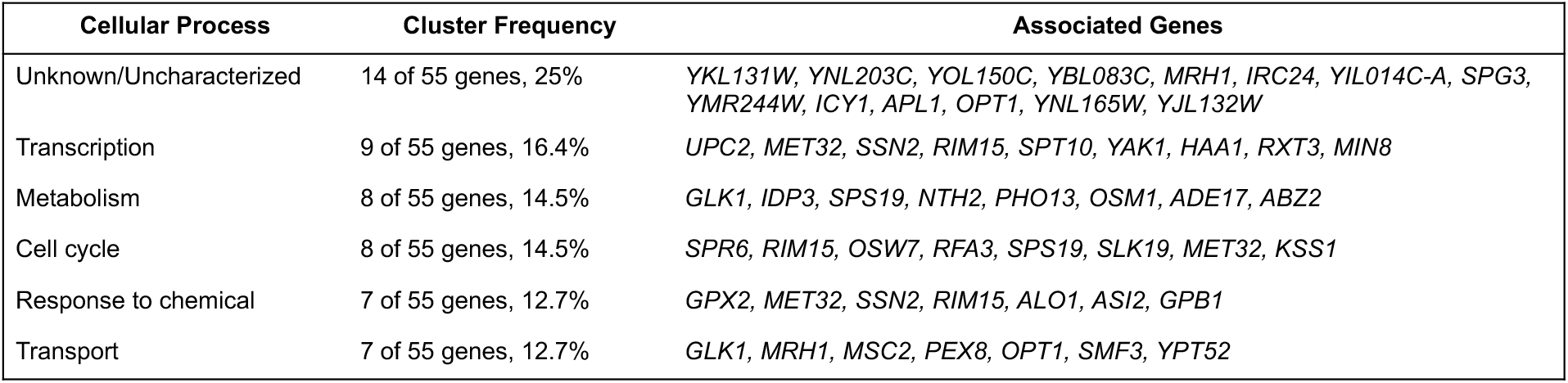
Clustering of identified gene deletions based on cellular process from a gene ontology (GO) cluster. Clusters from the GO were combined to create more broad groupings of cellular processes. For more information, please see *Materials and Methods*.

Once we identified the 55 unique strains, we conducted an initial evaluation of the ncAA incorporation efficiency of the identified strains by qualitatively evaluating their activities with analytical flow cytometry (SI Figure 2; this analysis included characterizations of multiple colonies identified as containing the same gene deletion). Based on these flow cytometry data and the barcode sequencing data, two gene deletions were determined to be of primary interest based on their presence in multiple sorting paths or because they were the dominant deletion strain in a sort track. One deleted gene, *YIL014C-A* (also known as *YIL015C-A*) which encodes a protein of unknown function, appeared in five different screens across both OTSs; this isolation in five different screens was the most of any candidate knockout. Colonies verified to contain this deletion exhibited consistently improved ncAA incorporation behavior compared to other strains (SI Figure 2). *YIL014C-A* encodes a protein with an uncharacterized function. Interestingly, the gene is located between genes encoding threonine and aspartate tRNAs. The deletion of *ALO1* appeared at high frequency in multiple sorting paths and exhibited improved ncAA incorporation compared to other deletion strains, as analyzed via flow cytometry (SI Figure 2). *ALO1* encodes for the enzyme D-arabino-1,4-lactone oxidase, which catalyzes the biosynthesis of a molecule that protects against oxidative stress (dehydro-D-arabinono-1,4-lactone) (*32*). Although *ALO1* plays a role in mitigating oxidative stress and is localized in the mitochondria (*33*), its deletion in another study was found to increase gross chromosomal rearrangements in yeast (*34*), suggesting that the gene and its protein product may be implicated in processes related to protein translation. While no direct connections of *ALO1* or *YIL014C-A* to protein biosynthesis are known, these two deletions were subjected to additional investigations to confirm their enhancement of ncAA incorporation efficiency in *S. cerevisiae*.

### Verifying improved ncAA incorporation efficiency of *yil014c-aΔ* and *alo1Δ*

To further verify the apparently enhanced ncAA incorporation discovered during post-FACS flow cytometry characterization, we used *yil014c-aΔ* and *alo1Δ* yeast strains from commercial sources. These independently acquired strains were transformed with one of two single plasmid system reporter/OTS plasmids: BXG-altTAG with either TyrOmeRS or LeuOmeRS (and corresponding tRNA_CUA_). Transformed knockout strains were quantitatively evaluated using two metrics: relative readthrough efficiency (RRE) and maximum misincorporation activity (MMF). These metrics were originally described by Barrick and coworkers in *E. coli* and extended to *S. cerevisiae* by our group (*27–29, 35, 36*). RRE measures ncAA incorporation efficiency, where a value of 1 indicates wildtype incorporation efficiency at the amber stop codon and a value of 0 indicates total translation truncation at the amber stop codon (SI Equation 1). MMF measures the extent of misincorporation of a canonical amino acid at the amber stop codon in the absences of ncAA. An MMF value of 0 indicates no detectable misincorporation, while values approaching 1 indicate high levels of canonical amino acid misincorporation (SI Equation 2) (*27–29, 35, 36*).

The availability of RRE and MMF evaluations allowed us to determine if improvements in ncAA incorporation efficiency due to a gene knockout would be conserved across different mating patterns. The YKO collection we screened consisted of heterozygous diploid yeast deletions (*S. cerevisiae* BY4743). Yeast BY4743 diploids are the product of haploid cells MAT**a** (BY4741) and MATα (BY4742). To our knowledge, no reports have characterized differences in ncAA incorporation efficiency between these three strains. To evaluate this question, we compared RRE and MMF in BY4741ΔPPQ1, BY4742ΔPPQ1, and BY4743ΔPPQ1 strains to the baseline incorporation efficiency in their respective parent strains. *ppq1Δ* was used because its enhanced ncAA incorporation efficiency has been previously demonstrated; in the absence of OTSs, deletion of the gene leads to a known phenotype of increased stop codon readthrough (*27, 37*). Using a two-plasmid reporter system that we have previously determined to be functionally equivalent to the single-plasmid systems used during screening (*29*), we cotransformed a plasmid encoding a BXG-altTAG reporter along with a plasmid encoding the TyrOmeRS/tRNA_CUA_^Tyr^ OTS into these six strains and evaluated their incorporation efficiency and fidelity with flow cytometry. Both RRE values and the flow cytometry dot plots suggest that enhanced ncAA incorporation due to deleting *PPQ1* are only observed in the MAT**a** (BY4741) and heterozygous diploid (BY4743) genetic backgrounds (SI Figure 3). Surprisingly, BY4742ΔPPQ1 exhibited decreased incorporation efficiency compared to BY4742. Based on these data, we limited further characterizations of incorporation efficiency to BY4741 and BY4743 strain backgrounds.

To characterize properties of strains possessing deletions of interest, we first wanted to verify that commercially acquired strains of *yil014c-aΔ* and *alo1Δ* possessed enhanced OmeY incorporation efficiency. We induced *yil014c-aΔ, alo1Δ*, and nonmutant parent cultures transformed with either LeuOmeRS-BXG-altTAG or TyrOmeRS-BXG-altTAG single plasmid systems in both BY4741 and BY4743 backgrounds with 1 mM OmeY and evaluated their incorporation efficiency and fidelity with flow cytometry. In cells expressing LeuOmeRS/tRNA_CUA_^Leu^ and induced with OmeY, BY4741ΔYIL014C-A and BY4741ΔALO1 strains possessed higher incorporation efficiency compared to BY4741, as demonstrated by higher RRE values. Since both gene deletion strains were identified in screens with the LeuOmeRS/tRNA_CUA_^Leu^ machinery, this result validates that these deletions improve ncAA incorporation efficiency (Figure 2A). Furthermore, the increased incorporation efficiency of the knockout strains does not appear to compromise the incorporation fidelity, as measured by the MMF metric; MMF values are the same or lower (improved) in comparison to the parent strain. Higher RRE values were observed with the TyrOmeRS/tRNA_CUA_^Tyr^ machinery in the BY4741 background with both deletions compared to the parent, and in the BY4743 background with ΔYIL014C-A (Figure 2A). The observation that *alo1Δ* improves ncAA incorporation efficiency with TyrOmeRS/tRNA_CUA_^Tyr^ is noteworthy because *alo1Δ* was only identified in screens utilizing LeuOmeRS/ tRNA_CUA_^Leu^. As determined by MMF measurements, ncAA incorporation fidelity in the knockout strains did not change or were improved compared to the parent strain. These data demonstrate that deleting *ALO1* and *YIL014C-A* from the yeast genome enhances ncAA incorporation efficiency while maintaining fidelity. Furthermore, the improvement of RRE with *alo1Δ* when utilizing TyrOmeRS suggests that deletion strains identified during screening can enhance incorporation efficiency using OTSs beyond the initial machinery used during discovery of candidate knockouts.

**Figure 2.**
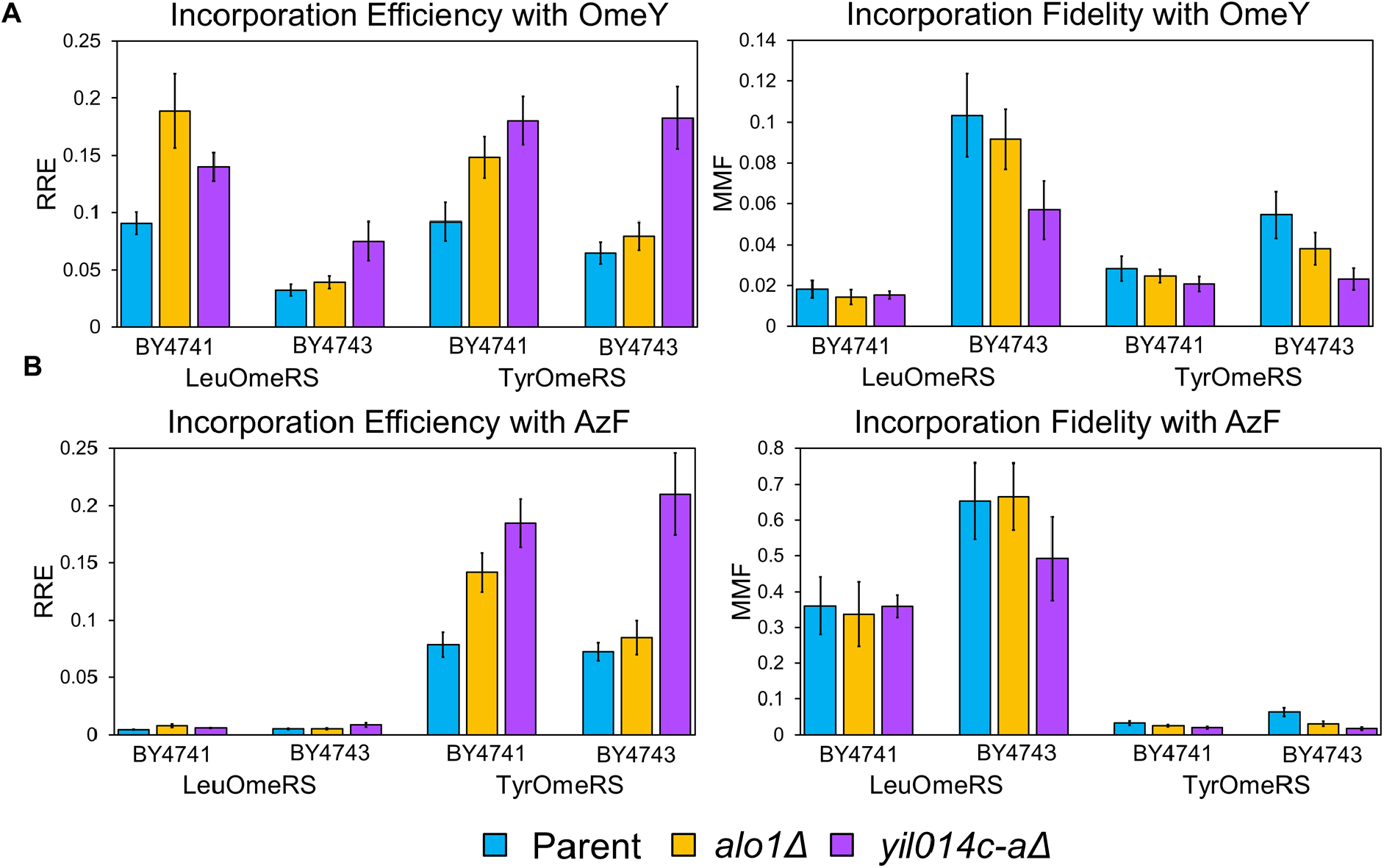
Quantitative evaluation of ncAA incorporation efficiency and fidelity with *yil014c-aΔ* and *alo1Δ* with OTSs used during screening. (A) RRE and MMF metrics based on a flow cytometry ncAA incorporation assay following the induction of cells in the presence of 1 mM OmeY. (B) RRE and MMF metrics based on flow cytometry ncAA incorporation assay following the induction of cells in the presence of 1 mM AzF (see Figure 3A for chemical structure). All samples in this figure were evaluated in biological triplicate. Error bars represent the standard deviation based on the propagated error. All experiments utilized BXG-altTAG single plasmid reporter constructs.

To further validate the enhanced incorporation efficiency of these strains, we induced the same cultures with *p*-azido-L-phenylalanine (AzF), a ncAA not utilized during screening. Prior work has shown that TyrOmeRS supports translational activity with AzF, while LeuOmeRS does not support AzF incorporation (*27*). In strains transformed with plasmids encoding the TyrOmeRS machinery, we observed similar improvements in incorporation efficiency with both knockout strains, with *yil014c-aΔ* and *alo1Δ* exhibiting higher RRE values (Figure 2B). We also found that, as expected, translational activity with the LeuOmeRS was not observed in either the parent strain or knockout strains. Consistent with experiments involving OmeY, the improvement in AzF incorporation efficiency did not appear to come at the expense of incorporation fidelity; MMF measurements made with both gene deletions exhibited values comparable to the MMF values determined with the parent strain. These characterizations verify that two leading candidate deletion strains identified via our screening campaign exhibit enhanced ncAA incorporation efficiency with a ncAA that was not used during the screening process. To further explore how these strains can be used in additional applications involving the preparation of ncAA-containing proteins, we extended our investigations to include a broader set of OTSs and ncAAs.

### Investigating activity with additional OTSs and ncAAs

We then sought to better understand if the enhanced incorporation efficiency observed in these deletion strains are dependent on the identity of the ncAAs and OTSs or if the strains promote more efficient ncAA incorporation in general. First, we evaluated the incorporation efficiency of *yil014c-aΔ* and *alo1Δ* strains with a OTS/ncAA combination not used during any portion of screening, SpecOPGRS-3. This OTS is an *E. coli* tyrosyl-tRNA synthetase variant that was engineered to incorporate *p*-propargyloxy-L-phenylalanine (OPG) while minimizing incorporation of several structurally related ncAAs include OmeY, AzF, *p*-acetyl-L-phenylalanine (AcF), *p*-propargyloxy-L-phenylalanine (OPG), 4-azidomethyl-L-phenylalanine (AzMF), and 4-iodo-L-phenylalanine (*15*). Three strains, BY4741, BY4741ΔALO1, and BY4741ΔYIL014C-A, were transformed with two separate plasmids encoding SpecOPGRS-3 and the BXG-altTAG reporter. These transformants were then induced in the presence of 1 mM OPG and their ncAA incorporation efficiency and fidelity was evaluated. Both *yil014c-aΔ* and *alo1Δ* mutant strains exhibited higher RRE values compared to the parent (Figure 3B), indicating that the knockout strains provide enhanced incorporation efficiencies when using SpecOPGRS-3 to encode OPG. To statistically demonstrate the increased incorporation efficiency of the two knockout strains, a one-way analysis of variance (ANOVA) statistical test was performed on the C-terminus (GFP) detections of the parent and two knockout strains. The statistical test reveals a significant increase in GFP levels in the two knockout strains over the parent strain, further confirming the improved ncAA incorporation efficiency of the two knockout strains in comparison to the parent strain (SI Figure 4).

**Figure 3.**
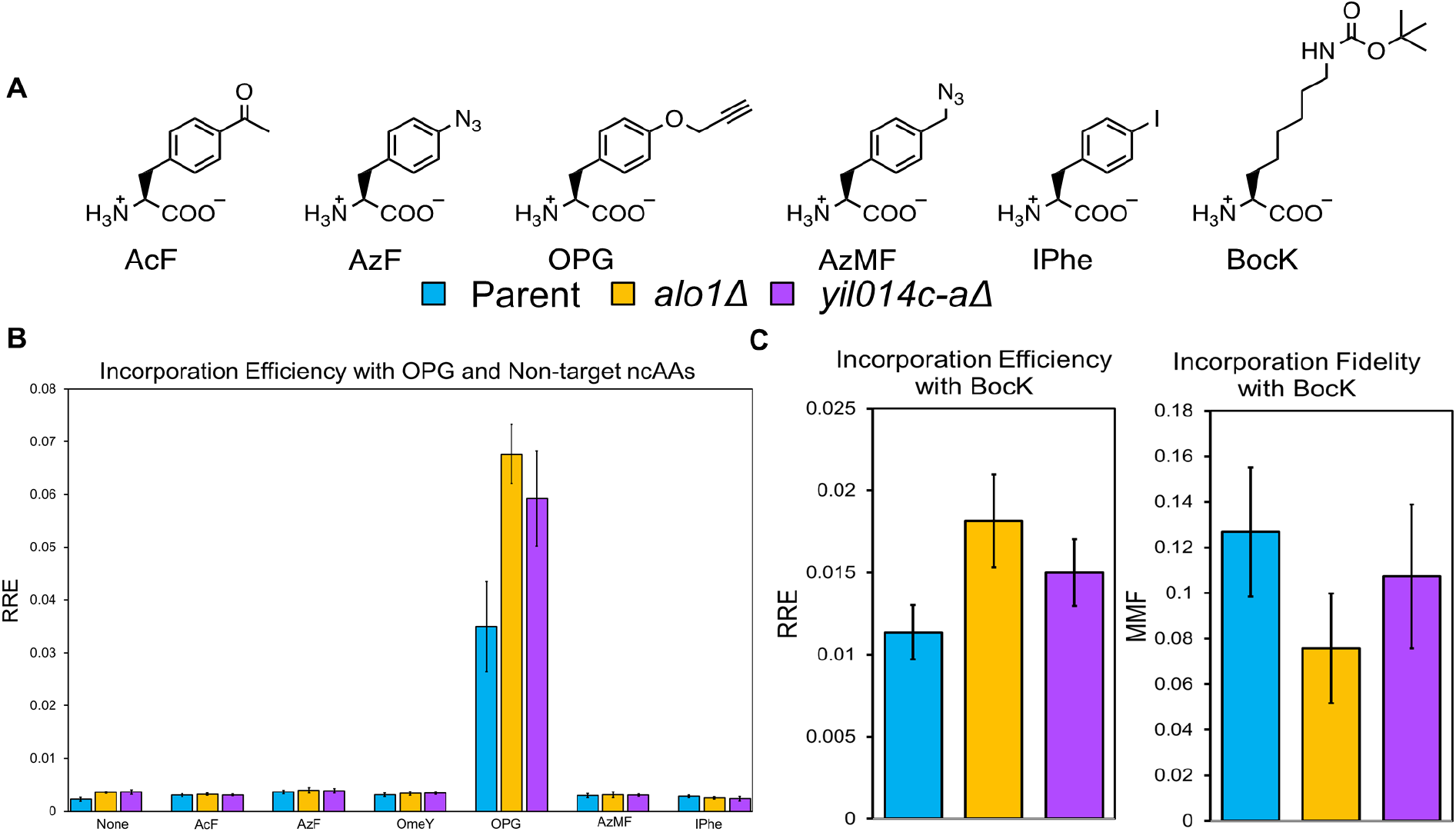
Quantitative evaluation of ncAA incorporation efficiency of *alo1Δ* and *yil014c-aΔ* with additional OTSs and ncAAs. (A) Structures of additional ncAAs used in these studies. (B) RRE metric based on a flow cytometry ncAA incorporation assay following the induction of cells expressing SpecOPGRS-3 in the absence of ncAA and in the presence of 1 mM OPG and five additional ncAAs. (C) RRE and MMF metric based on a flow cytometry ncAA incorporation assay following the induction of cells expressing a pyrrolysyl-tRNA synthetase/tRNA OTS in the presence of 10 mM BocK. Experiments shown in (B) and (C) utilize the BXG-altTAG fluorescent reporter cotransformed with the appropriate OTS on two separate plasmids. All samples were evaluated in biological triplicate. Error bars represent the standard deviation based on the propagated error.

Since we observed improved incorporation of OPG with ΔALO1 and ΔYIL014C-A, we wanted to determine if this improvement in OPG incorporation was accompanied by increases in the incorporation of the structurally similar “off-target” ncAAs. To evaluate the extent of specificity of the enhanced incorporation efficiency, the parent BY4741 and BY4741ΔALO1 and BY4741ΔYIL014C-A were induced in the presence of 1 mM of each of the five structurally similar ncAAs: OmeY, AzF, AcF, AzMF, and IPhe (Figure 3A, B). When induced in the presence of any of the five off-target ncAAs, no improvement in incorporation efficiency was observed with the two knockout strains compared to the parent strain as determined via RRE measurements (Figure 3B). In addition, MMF values suggest that there are comparable levels of canonical amino acid misincorporation between the parent and knockout strains, further demonstrating that the two knockout strains preserve protein translation fidelity. (SI Figure 5). The preservation of ncAA specificity by both knockout strains suggests that the deletions do not decrease the fidelity of protein translation.

To further explore the potential generality of ncAA incorporation enhancements conferred by *yil014c-aΔ* and *alo1Δ*, we investigated whether strains containing these deletions could enhance the activity of an OTS that is structurally and functionally distinct from the other OTSs used in this work. Recently, Van Deventer and coworkers demonstrated that a *Methanomethylophilus alvus* pyrrolysyl-tRNA synthetase (MaPylRS) can be utilized to encode ncAAs at amber codons in yeast (*38*). More broadly, PylRSs have supported the incorporation of hundreds of ncAAs relevant to drug development and biological studies in *E. coli* and mammalian cells (*39–42*), but their implementation in yeast remains limited. Strains cotransformed with two plasmids, MaPylRS/tRNA_CUA_^MaPyl^ and the BXG-altTAG reporter plasmid constructs, were induced with 10 mM *N_ε_*-Boc-L-lysine (BocK) (Figure 3A) and evaluated for ncAA incorporation efficiency and fidelity (*38*). We observed that BY4741ΔALO1 and BY4741ΔYIL014C-A exhibited improved incorporation efficiency of BocK compared to the parent BY4741 based on their respective RRE values (Figure 3C), despite the fact that the RRE values of all three strains are in the range of 1%. Qualitative analysis of the flow cytometry dot plots also shows increased GFP detection for both *yil014c-aΔ* and *alo1Δ* compared to the parent strain (SI Figure 6). One-way ANOVA tests of the MFI values of GFP fluorescence (C-terminus detection) also demonstrate a statistically significant increase in the two knockout strains compared to the parent strain (SI Figure 5). Consistent with the experiments conducted with other OTSs, the knockout strains maintained similar levels of incorporation fidelity compared to the parent strain. These findings demonstrate that the improved ncAA incorporation efficiencies conferred by deleting either *YIL014C-A* or *ALO1* are compatible with at least three types of OTSs and both aromatic and aliphatic ncAAs. Furthermore, the enhancements observed in a wide range of scenarios suggest that selective manipulations of the yeast genome have the potential to be advantageous for a wide range of applications requiring the efficient production of ncAA-containing proteins.

## Conclusion

In this work, we screened a pooled heterozygous diploid yeast knockout collection based on the phenotype of enhanced ncAA incorporation efficiency and identified 55 candidate single-gene knockout strains. Identified genes encode for proteins that participate in a wide range of cellular functions, many of which have no known connection to protein translation. We selected two promising candidate deletion strains, *yil014c-aΔ* and *alo1Δ*, and verified their improved behavior compared to parent strains in two mating backgrounds. Strains containing either deletion enhanced ncAA incorporation efficiency with a range OTSs and ncAAs, but without apparent loss of fidelity based on our reporter system-based evaluations. This observation holds for several OTSs, including a high-fidelity OTS that facilitates “on-target” ncAA incorporation of a single ncAA; the deletions do not compromise this fidelity. Finally, the enhancement of ncAA incorporation efficiency and fidelity we observed with OTSs derived from *E. coli* were confirmed when using the structurally distinct OTS based on the pyrrolysyl-tRNA synthetase from *Methanomethylophilus alvus*. While the focus of this study is on the development of the screening strategy and initial validation of the improved performance of knockouts, it will be important in future work to conduct mass spectrometry characterizations to gain further insights into ncAA incorporation efficiency and fidelity in strains containing the knockouts reported here. Overall, the improvement in ncAA incorporation efficiency we observed in these studies suggests that gene knockouts are a viable strategy for advancing genetic code manipulation in yeast.

This is the first study, to our knowledge, of a genome-wide screen in any organism focused on identifying processes that contribute to ncAA incorporation. Our findings reveal some of the complexities of the protein translation apparatus, and more specifically ncAA incorporation. While the screen we report here was conducted in *S. cerevisiae*, the broad range of genes identified in this work suggest that our screening approach is likely to be useful for discovering deletions that enhance genetic code manipulation in other organisms. The use of unbiased screens to elucidate key processes that affect ncAA incorporation has the potential to advance genetic code expansion as well as our fundamental understanding of protein biosynthesis. However, it remains to be determined to what extent insights gained from identifying processes that enhance ncAA incorporation apply to protein biosynthesis processes more broadly.

Comparison of the deletions we identified in this work with the deletions found in a prior genome-wide screen in *S. cerevisiae* focused on premature stop codon bypass (without the use of OTSs or ncAAs) indicates that identified hits are generally nonoverlapping between the two studies (*43*). In addition, one of the key gene deletions we investigated here, *YIL014C-A*, does not have a known homolog in mammalian cells, suggesting that some screening hits will be species-specific. In any case, our report provides an important step towards engineering *S. cerevisiae* to accommodate expanded genetic codes. Further studies to evaluate incorporation efficiency with additional candidate deletion strains and in multi-gene knockout strains present relatively straightforward ways to extend the current work. In addition, our framework for discovering and evaluating candidate strains will facilitate studies comparing ncAA incorporation in yeast of different mating types and genetic backgrounds to elucidate why single-gene deletions that enhance ncAA incorporation efficiency in one strain background may not result in enhanced efficiency in others. There are many important questions to be answered at the interface of synthetic and fundamental biology to facilitate effective enhancement of genetic code manipulation.

Larger genome engineering strategies are important tools for constructing cells with improved abilities to accommodate expanded genetic codes, and numerous resources in *S. cerevisiae* may facilitate implementation of such strategies. The ongoing Synthetic Yeast Genome Project (Sc2.0) seeks to elaborate genomic design and construction strategies, partially with an eye on enhancing ncAA incorporation via the removal of the amber (TAG) stop codon from the yeast genome (*19, 20, 44, 45*). The Sc2.0 project also facilitates the genomic diversification and screening of yeast strains via Synthetic Chromosome Rearrangement and Modification by LoxPsym-mediated Evolution (SCRaMbLE) (*46, 47*). This approach has been shown to support discovery of strains exhibiting several improved phenotypes and may also facilitate enhancement of ncAA incorporation efficiency. In addition to the tools of the Sc2.0 project, several CRISPR/Cas-based approaches in *S. cerevisiae* are suitable for performing genomic edits in high throughout (*48–50*). Furthermore, both CRISPR/Cas systems and more “classical” genetic tools have been shown to be effective for modulating gene expression with an eye on improving yields of metabolic biosynthesis (*51–53*). Engineering efficient cellular syntheses of complex small molecules has many parallels to manipulating the processes required to enhance protein biosynthesis to function better with expanded genetic codes. Leveraging these extensive tools has the potential to establish well-defined strategies for enhancing ncAA incorporation while also yielding fundamental insights regarding protein translation. Our results here indicate that there are numerous cellular components beyond the known elements of the protein translation apparatus that can play important roles in enhancing ncAA incorporation efficiency. Our findings present a framework for exploring the biology that facilitates ncAA incorporation and presents strategies to improve the accommodation of expanded genetic codes in cells.

## Materials and Methods

### Materials

Synthetic oligonucleotides for PCR amplification and sequencing were purchased from GENEWIZ. Sanger sequencing was performed by Quintara Biosciences in Cambridge, Massachusetts. Epoch Life Science GenCatch Plasmid DNA Mini-Prep Kits were used for DNA purification from *E. coli* and for post-PCR cleanup for the barcode sequence analysis. Zymo Research Frozen-EZ Yeast Transformation II kits were used to make chemically competent yeast strains and to perform plasmid transformation in yeast. NcAAs were purchased from these companies: *p*-acetyl-L-phenylalanine (SynChem); *p*-azido-L-phenylalanine (Chem-Impex International); *O*-methyl-L-tyrosine (Chem-Impex International); *p*-propargyloxy-L-phenylalanine (Iris Biotech); 4-azidomethyl-L-phenylalanine (SynChem); 4-iodo-L-phenylalanine (AstaTech); *N_ε_*-Boc-L-lysine (Chem-Impex International).

### Media preparation and yeast strain construction

Liquid and solid media were prepared as previously reported (*28*). SD-SCAA and SG-SCAA media were prepared without leucine and uracil. Media with leucine and uracil dropped out were used for cells transformed with a dual-plasmid system (SI Table 1). 100x leucine was supplemented to the media at a final concentration of 1x leucine (*28*) for selection of cells transformed with single-plasmid systems. The molecular barcoded heterozygous diploid BY4743 yeast knockout collection was a gift from the laboratory of Maitreya Dunham at the University of Washington (*26*). BY4741ΔALO1 and BY4743ΔALO1 were purchased from Horizon Discovery. BY4741ΔYIL014C-A and BY4743ΔYIL014C-A were purchased from Euroscarf. The BY4742 knockout strains were a gift from the laboratory of Catherine Freudenreich at Tufts University.

### Preparing ncAA liquid stocks

Liquid stocks of ncAAs were prepared at a concentration of 50 mM L-isomer. Approximately 80% of the total volume of DI water was added to the solid ncAA. Following mixing, 6.0 N NaOH was added gradually to dissolve the ncAA. Following additional vortexing and complete dissolving of the ncAA, additional DI water was added to achieve the final volume. Stocks were sterile filtered with 0.2 μm filters. For OmeY, the solution was adjusted to pH 7 before filtering.

### YKO library electroporation and characterization

4 μg pRS416-BXG-TyrOmeRS, pRS416-BXGAltTAG-LeuOmeRS, and pRS416-BXGAltTAG-TyrOmeRS and 8 μg pRS416-BXG-LeuOmeRS were concentrated using Pellet Paint® NF Co-precipitant based on manufacturer’s protocols (*15*). Electrocompetent pooled YKO cell preparation and electroporations were followed as previously described (*54*), with YPD growth medium supplemented with 200 μg/mL G418 during propagation of YKO cells. To determine the transformation efficiency, electroporated cells were plated on SD-SCAA (–TRP –LEU –URA), and the remainder of cells were recovered in 100 mL SD-SCAA (–URA) supplemented with G418 (We note that G418 is not active in minimal media; however, to the best of our knowledge the presence of G418 during initial library growth steps did not appear to overtly interfere with the discovery of strains of interest). One electroporation was performed for each reporter/OTS plasmid. Plated dilutions were grown at 30 °C for 3–4 days and colonies were counted to calculate the number of individual transformants. The library recovery cultures were grown at 30 °C with shaking at 300 rpm to saturation, then expanded into 1 L SD-SCAA (–URA, 100 μg/mL G418) and grown at 30 °C with shaking overnight. The cells from each 1 L culture were pelleted and resuspended in 60% glycerol to result in a final concentration of 15% glycerol, aliquoted to cryogenic vials, and stored at −80 °C. At least 7 × 10^9^ cells were stored per vial. A 5 mL SD-SCAA (–URA) culture of each library was passaged in parallel to characterize the unsorted libraries using flow cytometry and to isolate individual colonies and conduct sequencing to determine YKO barcodes.

### YKO barcode sequencing

Fresh agar plates were streaked from either glycerol stocks or from liquid cultures. Isolated yeast colonies from agar plates were inoculated in 2 mL YPD cultures supplemented with penicillin-streptomycin overnight. Samples were then centrifuged in microcentrifuge tubes and the supernatant removed. To extract the genomic DNA, samples were resuspended in 200 μL 200 mM lithium acetate 1% SDS solution and incubated for 15 min then spun down and aspirated. Two washes, one with 600 μL 100% ethanol and one with 500 μL 70% ethanol, were performed before resuspending the DNA pellet in 600 μL sterile water. Next, PCR was performed on the genomic DNA to amplify the unique molecular barcodes. 1 μL genomic DNA was added to 1 μL 2% DMSO, 2.5 μL each 10 μM forward primer (5’-GATGTCCACGAGGTCTCT-3’) and 10 μM reverse primer (5’-CGGTGTCGGTCTCGTAG-3’), 25 μL 2X Q5 Master Mix, and 18 μL sterile water. Samples were incubated in a thermal cycler at 98°C for 30 s followed by 30 cycles at 98°C for 10 s, 58°C for 30 s, and 72°C for 30 s. Following these cycles, samples underwent a final extension at 72°C for 2 min. PCR cleanup was performed with an Epoch Life Science GenCatch Gel Extraction Kit. 450 μL GEX buffer was added to 45 μL of PCR product in a miniprep column, then spun down and the flow through discarded. 700 μL of WS was added to the column, and the sample was centrifuged. The column was then transferred to an empty microcentrifuge tube, and the sample was eluted with 40 μL of sterile water. Following a 3 min incubation, samples were centrifuged again, and the eluted DNA was prepared for Sanger sequencing. For each sample, two Sanger sequencing samples were prepared, one for sequencing the UPTAG and the second for the DNTAG. 10 μL DNA at 10 ng/μL was added to a PCR tube, along with 5 μL of 5 μM of either forward (5’-CGCCTCGACATCATCTGCCCAGATGCG-3’) or reverse (5’-GGATGTATGGGCTAAATGTACGGGGCG-3’) (*26*). Samples were then submitted for Sanger sequencing. To analyze the data, sequences were aligned with a template deletion cassette module, which revealed the 20mer UPTAG and DNTAG barcodes.

### Yeast transformations, propagation, and induction

Transformations were performed on Zymo competent *S. cerevisiae* BY4741, BY4742, BY4743 or single-gene deletions. For verification experiments utilizing single plasmids (i.e., pRS416-BXG-altTAG-LeuOmeRS or pRS416-BXG-altTAG-TyrOmeRS), cells were plated on solid SD-SCAA media (-URA) and grown at 30°C until colonies appeared (2–4 days). For verification experiments using two-plasmid systems, OTS-containing plasmids (pRS315-KanRmod-MaPylRS or pRS315-SpecOPGRS-3) were cotransformed with reporter plasmid pRS416-BXG-altTAG on solid SD-SCAA media (–LEU – URA). Cells were grown at 30°C until colonies appeared. Once colonies had grown, they were inoculated in 5 mL SD-SCAA media (with the same dropout conditions as the solid plates) in biological triplicate. Penicillin-streptomycin was added to all propagation and induction cultures to prevent bacterial transformation. The 5 mL propagation cultures were grown at 30°C for 2 days with shaking at 300 rpm. The day prior to flow cytometry, cultures were diluted in 5 mL SD media at OD_600_ = 1.4–8 hr following dilutions, cultures were induced in 2 mL SG-SCAA media (with the same dropout conditions as before) at OD_600_ = 1. Each biological replicate was induced both in the absence and presence of ncAA. Cultures were grown at 20 °C for 16 hr shaking at 300 rpm.

### Flow cytometry data collection and analysis

Induced samples were aliquoted in 96-well V-bottom plates. Detailed protocols describing the preparation of cultures for flow cytometry were reported in detail previously (*27, 28*). Analytical flow cytometry was performed on an Attune NxT (Life Technologies) at the Science and Technology Center, Tufts University. Analysis of flow cytometry data was performed with FlowJo and Microsoft Excel.

### Fluorescence-activated cell sorting (FACS)

Following induction, 30 million cells from each of the transformed yeast knockout libraries were pelleted and washed thrice with PBSA. Cells were suspended in PBSA prior to loading in the sorter. A S3e cell sorter (Bio-Rad) at the Science and Technology Center, Tufts University was used for sorting experiments. Sorting gates were drawn to acquire the top ~0.05% of cells with the greatest GFP fluorescence levels using the FL1 and FL2 detection channels on the instrument (this sorter lacks a laser suitable for BFP excitation). FL1 channel: 525/30 nm bandpass filter that captures main portion of GFP fluorescence emission. FL2 channel: 586/25 nm bandpass filter that captures high wavelength tail of GFP fluorescence emission. Collected events were deposited in 14 mL culture tubes containing 1 mL SD-SCAA (–URA) media. Sorting continued until 1,000 events were collected. Following sorting, the recovery media was supplemented to 5 mL and grown at 30 °C for 2 days.

### Gene ontology (GO) term mapping and groupings

All identified genes were grouped by cellular process using the Gene Ontology Slim Term Mapper. For Table 2, groupings of genes by broader categories were compiled by combining genes belonging to the following GO terms: Transcription: transcription by RNA polymerase I, transcription by RNA polymerase II; Metabolism: nucleobase-containing small molecule process, generation of precursor metabolites and energy; Cell Cycle: meiotic cell cycle, mitotic cell cycle, regulation of cell cycle; Transport: transmembrane transport, carbohydrate transport, endocytosis, Golgi vesicle transport, vesicle organization, ion transport, exocytosis; Response to chemical: response to chemical.

## Acknowledgements

This research was supported by a grant from the National Institute of General Medical Sciences of the National Institutes of Health (1R35GM133471). M.T.Z. was supported in part by the Nathan Gantcher Student Summer Scholars Program at Tufts University. J.T.S. was supported in part by a National Science Foundation Graduate Research Fellowship (ID: 2016231237). The content of this work is solely the responsibility of the authors and does not necessarily represent the official views of the National Institutes of Health or the National Science Foundation. The authors would like to thank Maitreya Dunham (University of Washington) for the pooled yeast knockout collection and Catherine Freudenreich (Tufts University) for BY4742 knockout strains. The authors would also like to thank Arlinda Rezhdo and Manjie Huang for their support in using the S3e cell sorter.

## Author Contributions

Conceptualization, M.T.Z, J.T.S., and J.A.V.; methodology, M.T.Z., J.T.S., and J.A.V.; investigation, M.T.Z. and J.T.S.; writing, M.T.Z.; review and editing, M.T.Z., J.T.S., and J.A.V.; funding acquisition, J.A.V.

## Supporting Information

The supporting information contains comprehensive tables detailing screening paths, plasmids used, and gene deletions identified. Additionally, figures containing flow cytometry plots and RRE and MMF data for certain experiments are also included.

## Author Information

### Corresponding Author

**James A. Van Deventer** – *Chemical and Biological Engineering Department, Tufts University, Medford, Massachusetts 02155, United States; Biomedical Engineering Department, Tufts University, Medford, Massachusetts 02155, United States*;

### Authors

**Matthew T. Zackin** – *Chemical and Biological Engineering Department, Tufts University, Medford, Massachusetts 02155, United States*;

**Jessica T. Stieglitz** – *Chemical and Biological Engineering Department, Tufts University, Medford, Massachusetts 02155, United States*;

## Declaration of Interests

Authors J.T.S. and J.A.V. have submitted a patent application for the aaRS sequence of SpecOPGRS-3: U.S. Provisional Application No. 63/190,336.

## References

1. J. W. Chin, Expanding and reprogramming the genetic code. Nature 550, 53–60 (2017).

2. K. Lang, J. W. Chin, Bioorthogonal reactions for labeling proteins. ACS Chem Biol 9, 16–20 (2014).

3. D. C. Dieterich, A. J. Link, J. Graumann, D. A. Tirrell, E. M. Schuman, Selective identification of newly synthesized proteins in mammalian cells using bioorthogonal noncanonical amino acid tagging (BONCAT). P Natl Acad Sci USA 103, 9482–9487 (2006).

4. A. Chatterjee, J. Guo, H. S. Lee, P. G. Schultz, A genetically encoded fluorescent probe in mammalian cells. J Am Chem Soc 135, 12540–12543 (2013).

5. J. Luo et al., Genetically encoded optochemical probes for simultaneous fluorescence reporting and light activation of protein function with two-photon excitation. J Am Chem Soc 136, 15551–15558 (2014).

6. H. S. Park et al., Expanding the genetic code of Escherichia coli with phosphoserine. Science 333, 1151–1154 (2011).

7. C. Hoppmann et al., Site-specific incorporation of phosphotyrosine using an expanded genetic code. Nat Chem Biol 13, 842–+ (2017).

8. X. Z. Luo et al., Genetically encoding phosphotyrosine and its nonhydrolyzable analog in bacteria. Nat Chem Biol 13, 845–+ (2017).

9. F. Tian et al., A general approach to site-specific antibody drug conjugates. P Natl Acad Sci USA 111, 1766–1771 (2014).

10. A. Rezhdo, M. Islam, M. J. Huang, J. A. Van Deventer, Future prospects for noncanonical amino acids in biological therapeutics. Curr Opin Biotech 60, 168–178 (2019).

11. D. J. Mandell et al., Biocontainment of genetically modified organisms by synthetic protein design (vol 518, pg 55, 2015). Nature 527, 264–264 (2015).

12. K. H. Wang, H. Neumann, S. Y. Peak-Chew, J. W. Chin, Evolved orthogonal ribosomes enhance the efficiency of synthetic genetic code expansion. Nat Biotechnol 25, 770–777 (2007).

13. B. J. Des Soye, J. R. Patel, F. J. Isaacs, M. C. Jewett, Repurposing the translation apparatus for synthetic biology. Curr Opin Chem Biol 28, 83–90 (2015).

14. D. B. F. Johnson et al., RF1 knockout allows ribosomal incorporation of unnatural amino acids at multiple sites. Nat Chem Biol 7, 779–786 (2011).

15. J. T. Stieglitz, and Van Deventer, J. A., High-throughput aminoacyl-tRNA synthetase engineering for genetic code expansion in yeast. bioRxiv, (2021).

16. O. Vargas-Rodriguez, A. Sevostyanova, D. Soll, A. Crnkovic, Upgrading aminoacyl-tRNA synthetases for genetic code expansion. Curr Opin Chem Biol 46, 115–122 (2018).

17. N. M. Reynolds, O. Vargas-Rodriguez, D. Soll, A. Crnkovic, The central role of tRNA in genetic code expansion. Biochim Biophys Acta Gen Subj 1861, 3001–3008 (2017).

18. C. C. Liu, P. G. Schultz, Adding New Chemistries to the Genetic Code. Annual Review of Biochemistry, Vol 79 79, 413–444 (2010).

19. N. Ostrov et al., Design, synthesis, and testing toward a 57-codon genome. Science 353, 819–822 (2016).

20. J. Fredens et al., Total synthesis of Escherichia coli with a recoded genome. Nature 569, 514–+ (2019).

21. J. D. Keasling, Manufacturing Molecules Through Metabolic Engineering. Science 330, 1355–1358 (2010).

22. G. Giaever et al., Functional profiling of the Saccharomyces cerevisiae genome. Nature 418, 387–391 (2002).

23. G. Giaever, C. Nislow, The Yeast Deletion Collection: A Decade of Functional Genomics. Genetics 197, 451–465 (2014).

24. N. P. Mira, M. C. Teixeira, I. Sa-Correia, Adaptive Response and Tolerance to Weak Acids in Saccharomyces cerevisiae: A Genome-Wide View. Omics 14, 525–540 (2010).

25. A. Sliva, Z. Kuang, P. B. Meluh, J. D. Boeke, Barcode Sequencing Screen Identifies SUB1 as a Regulator of Yeast Pheromone Inducible Genes. G3-Genes Genom Genet 6, 881–892 (2016).

26. X. W. Pan et al., A robust toolkit for functional profiling of the yeast genome. Mol Cell 16, 487–496 (2004).

27. K. A. Potts, J. T. Stieglitz, M. Lei, J. A. Van Deventer, Reporter system architecture affects measurements of noncanonical amino acid incorporation efficiency and fidelity. Mol Syst Des Eng 5, 573–588 (2020).

28. J. T. Stieglitz, H. P. Kehoe, M. Lei, J. A. Van Deventer, A Robust and Quantitative Reporter System To Evaluate Noncanonical Amino Acid Incorporation in Yeast. Acs Synth Biol 7, 2256–2269 (2018).

29. J. T. Stieglitz, K. A. Potts, J. A. Van Deventer, Broadening the Toolkit for Quantitatively Evaluating Noncanonical Amino Acid Incorporation in Yeast. Acs Synth Biol 10, 3094–3104 (2021).

30. C. Payen et al., High-Throughput Identification of Adaptive Mutations in Experimentally Evolved Yeast Populations. Plos Genet 12, (2016).

31. J. M. Cherry et al., Saccharomyces Genome Database: the genomics resource of budding yeast. Nucleic Acids Res 40, D700–D705 (2012).

32. W. K. Huh et al., D-Erythroascorbic acid is an important antioxidant molecule in Saccharomyces cerevisiae. Mol Microbiol 30, 895–903 (1998).

33. A. Sickmann et al., The proteome of Saccharomyces cerevisiae mitochondria. P Natl Acad Sci USA 100, 13207–13212 (2003).

34. S. Smith et al., Mutator genes for suppression of gross chromosomal rearrangements identified by a genome-wide screening in Saccharomyces cerevisiae. P Natl Acad Sci USA 101, 9039–9044 (2004).

35. J. W. Monk et al., Rapid and Inexpensive Evaluation of Nonstandard Amino Acid Incorporation in Escherichia coli. Acs Synth Biol 6, 45–54 (2017).

36. M. Islam et al., Chemical Diversification of Simple Synthetic Antibodies. Acs Chemical Biology 16, 344–359 (2021).

37. T. von der Haar, M. F. Tuite, Regulated translational bypass of stop codons in yeast. Trends Microbiol 15, 78–86 (2007).

38. J. T. Stieglitz, P. Lahiri, M. I. Stout, J. A. Van Deventer, Exploration of Methanomethylophilus alvus Pyrrolysyl-tRNA Synthetase Activity in Yeast. Acs Synth Biol, (2022).

39. W. Wan, J. M. Tharp, W. R. Liu, Pyrrolysyl-tRNA synthetase: An ordinary enzyme but an outstanding genetic code expansion tool. Bba-Proteins Proteom 1844, 1059–1070 (2014).

40. J. C. W. Willis, J. W. Chin, Mutually orthogonal pyrrolysyl-tRNA synthetase/tRNA pairs. Nat Chem 10, 831–837 (2018).

41. V. Beranek, J. C. W. Willis, J. W. Chin, An Evolved Methanomethylophilus alvus Pyrrolysyl-tRNA Synthetase/tRNA Pair Is Highly Active and Orthogonal in Mammalian Cells. Biochemistry-Us 58, 387–390 (2019).

42. I. Coin, Application of non-canonical crosslinking amino acids to study proteinprotein interactions in live cells. Curr Opin Chem Biol 46, 156–163 (2018).

43. B. Samanfar et al., A global investigation of gene deletion strains that affect premature stop codon bypass in yeast, Saccharomyces cerevisiae. Mol Biosyst 10, 916–924 (2014).

44. J. Sanders, S. A. Hoffmann, A. P. Green, Y. Cai, New opportunities for genetic code expansion in synthetic yeast. Curr Opin Biotechnol 75, 102691 (2022).

45. S. M. Richardson et al., Design of a synthetic yeast genome. Science 355, 1040–1044 (2017).

46. J. Y. Ong et al., SCRaMbLE: A Study of Its Robustness and Challenges through Enhancement of Hygromycin B Resistance in a Semi-Synthetic Yeast. Bioengineering-Basel 8, (2021).

47. W. Liu et al., Rapid pathway prototyping and engineering using in vitro and in vivo synthetic genome SCRaMbLE-in methods. Nat Commun 9, (2018).

48. Z. H. Bao et al., Genome-scale engineering of Saccharomyces cerevisiae with single-nucleotide precision. Nat Biotechnol 36, 505–+ (2018).

49. X. G. Guo et al., High-throughput creation and functional profiling of DNA sequence variant libraries using CRISPR-Cas9 in yeast. Nat Biotechnol 36, 540–+ (2018).

50. K. R. Roy et al., Multiplexed precision genome editing with trackable genomic barcodes in yeast. Nat Biotechnol 36, 512–+ (2018).

51. E. K. Bowman et al., Bidirectional titration of yeast gene expression using a pooled CRISPR guide RNA approach. P Natl Acad Sci USA 117, 18424–18430 (2020).

52. G. M. Jones et al., A systematic library for comprehensive overexpression screens in Saccharomyces cerevisiae. Nat Methods 5, 239–241 (2008).

53. X. S. Xu, L. S. Oi, A CRISPR-dCas Toolbox for Genetic Engineering and Synthetic Biology. J Mol Biol 431, 34–47 (2019).

54. J. A. Van Deventer, K. D. Wittrup, Yeast Surface Display for Antibody Isolation: Library Construction, Library Screening, and Affinity Maturation. Methods Mol Biol 1131, 151–181 (2014).

