## Supporting Information for "Genome-wide screen for enhanced noncanonical amino acid incorporation in yeast"

**Table S1.** Plasmids utilized in this study.

| Plasmid 1 | Plasmid 2 | Purpose |
|---|---|---|
| pRS416-BXG-altTAG-LeuOmeRS | N/A | Screening, incorporation assay |
| pRS416-BXG-altTAG-TyrOmeRS | N/A | Screening, incorporation assay |
| pRS416-BXG-LeuOmeRS | N/A | Screening |
| pRS416-BXG-TyrOmeRS | N/A | Screening |
| pRS416-BYG-LeuOmeRS | N/A | WT reporter |
| pRS416-BYG-TyrOmeRS | N/A | WT reporter |
| pRS416-BXG-altTAG | pRS315-TyrOmeRS | Incorporation assay |
| pRS416-BYG | pRS315-TyrOmeRS | WT reporter |
| pRS416-BXG-altTAG | pRS315-SpecOPGRS-3 | Incorporation assay |
| pRS416-BYG | pRS315-SpecOPGRS-3 | WT reporter |
| pRS416-BXG-altTAG | pRS315-KanRmod-MaPylRS | Incorporation assay |
| pRS416-BYG | pRS315-KanRmod-MaPylRS | WT reporter |

**Table S2.** Transformation efficiency of the four transformed YKO libraries.

| | |
|---|---|
| LeuOmeRS-BXG-altTAG | $3.0 \times 10^4$ transformants |
| LeuOmeRS-BXG | $1.3 \times 10^5$ transformants |
| TyrOmeRS-BXG-altTAG | $2.6 \times 10^5$ transformants |
| TyrOmeRS-BXG | $1.9 \times 10^5$ transformants |

**Table S3.** Initial barcode sequence characterizations for the four naïve libraries to evaluate strain diversity prior to screening.

| TyrBXG | TyrAlt | LeuBXG | LeuAlt |
|---|---|---|---|
| YCR036W | YLR070C | YIL121W | YJR005W |
| YNL009W | YIR014W | YOR339C | YPL020C |
| YMR161W | YHR033W | YER087C | YBL105C |
| YJR075W | YPR035W | YIL003W | YEL063C |
| YLR320W | YLL048C | YIL011W | YOL138C |
| YOR043W | YDR232W | YLR382C | YDL152W |
| YIL137C | YIL121W | YPR165W | YDL152W |
| YMR285C | YBR033W | YPR165W | YGR216C |
| | | YDL129W | YGL047W |
| | | | YGR209C |
| 8 unique<br>8 sequenced | 8 unique<br>8 sequenced | 8 unique<br>9 sequenced | 9 unique<br>10 sequenced |

**Table S4.** Overview of the screening conditions for the 12 sorting paths performed on a Biorad s3e cell sorter. FACS used FL1, FL2 detection channels to isolate cells exhibiting high levels of GFP (this instrumentation lacks a laser suitable for BFP excitation). FL1 channel: 525/30 nm bandpass filter that captures main portion of GFP fluorescence emission. FL2 channel: 586/25 nm bandpass filter that captures high wavelength tail of GFP fluorescence emission.

| Screening Path | Synthetase | Reporter | OmeY Induction Concentration | FACS Detector |
|---|---|---|---|---|
| 488_0 | LeuOmeRS | BXG-altTAG | 0 | FL1 |
| 488_0.1 | LeuOmeRS | BXG-altTAG | 0.1 mM | FL1 |
| 488_1 | LeuOmeRS | BXG-altTAG | 1 mM | FL1 |
| LA0 | LeuOmeRS | BXG-altTAG | 0 | FL1, FL2 |
| LA0.1 | LeuOmeRS | BXG-altTAG | 0.1 mM | FL1, FL2 |
| LA1 | LeuOmeRS | BXG-altTAG | 1 mM | FL1, FL2 |
| LX0.1 | LeuOmeRS | BXG | 0.1 mM | FL1, FL2 |
| LX1 | LeuOmeRS | BXG | 1 mM | FL1, FL2 |
| YX0.1 | TyrOmeRS | BXG | 0.1 mM | FL1, FL2 |
| YX1 | TyrOmeRS | BXG | 1 mM | FL1, FL2 |
| YA0.1 | TyrOmeRS | BXG-altTAG | 0.1 mM | FL1, FL2 |
| YA1 | TyrOmeRS | BXG-altTAG | 1 mM | FL1, FL2 |

**Table S5.** Comprehensive list of identified strains. Gene descriptions were compiled from the results of a Gene Ontology (GO) search using the *Saccharomyces* Genome Database (SGD).

| Gene | Simplified Name | Gene Ontology Cellular Process Term | Sort(s) Identified | Frequency | Total |
|---|---|---|---|---|---|
| YJL173C | RFA3 | Meiotic cell cycle, protein modification by small protein conjugation or removal, DNA repair, organelle fission, telomere organization, DNA recombination, DNA replication | LA0 | 5 | 5 |
| YIL015C-A | N/A | No annotated term | LA0, LA0.1, LA1, 488_0, YA1.0 | 2 + 2 + 8 + 5 + 1 | 18 |
| YML086C | ALO1 | Response to chemical, response to oxidative stress | LA0.1, LA1 | 2 + 1 | 3 |
| YPR116W | RRG8 | tRNA processing, mitochondrion organization | LA0 | 2 | 2 |
| YOR195W | SLK19 | Meiotic cell cycle, regulation of cell cycle, cytoskeleton organization, mitotic cell cycle, organelle fission, chromosome segregation | LA0.1 | 1 | 1 |
| YHL034C | SBP1 | Regulation of translation | LA1 | 1 | 1 |
| YGR249W | MGA1 | Pseudohyphal growth | 488_0, YX0.1 | 1 + 1 | 2 |
| YKL131W | N/A | No annotated term | LA0.1, 488_0 | 3 + 2 | 5 |
| YMR289W | ABZ2 | Vitamin metabolic stress | 488_0.1 | 9 | 9 |
| YOR371C | GPB1 | Response to chemical, protein modification by small protein conjugation or removal, proteolysis involved in cellular protein catabolic process | 488_1, YA0.1 | 2 + 1 | 3 |
| YJR005W | APL1 | Pseudohyphal growth, invasive growth in response to glucose limitation | 488_1, YA0.1 | 1 + 2 | 3 |
| YMR244W | N/A | No annotated term | 488_1 | 1 | 1 |
| YDL076C | RXT3 | Transcription by RNA polymerase I | 488_1 | 1 | 1 |
| YDR504C | SPG3 | No annotated term | 488_1 | 1 | 1 |
| YJL127C | SPT10 | Transcription by RNA polymerase II, DNA repair, telomere organization, chromatin organization, peptidyl-amino acid modification, nucleus organization, protein acylation, histone modification | YX0.1 | 1 | 1 |
| YNL090W | RHO2 | Cytoskeleton organization, cell wall organization or biogenesis | YX0.1 | 1 | 1 |
| YMR195W | ICY1 | No annotated term | YX0.1 | 1 | 1 |
| YPR008W | HAA1 | Transcription by RNA polymerase II | YX0.1 | 1 | 1 |
| YBR001C | NTH2 | Oligosaccharide metabolic process | YX0.1 | 1 | 1 |
| YDR253C | MET32 | Transcription by RNA polymerase II, regulation of cell cycle, mitotic cell cycle | YX0.1 | 1 | 1 |
| YDR443C | SSN2 | Response to chemical, transcription by RNA polymerase II | YX0.1 | 1 | 1 |
| YJL132W | N/A | No annotated term | YX0.1 | 1 | 1 |
| YMR120C | ADE17 | Nucleobase-containing small molecule metabolic process | LX1.0 | 1 | 1 |
| YKR014C | YPT52 | Endocytosis, protein targeting, organelle assembly, endosomal transport, Golgi vesicle transport, vesicle organization | LX1.0 | 1 | 1 |
| YPR010C | RPA135 | Transcription by RNA polymerase I | LX1.0 | 1 | 1 |
| YNL203C | N/A | No annotated term | LX1.0 | 2 | 2 |
| YNL159C | ASI2 | Response to chemical, proteolysis involved in cellular protein catabolic process | LX1.0 | 1 | 1 |
| YLR034C | SMF3 | Ion transport, cellular ion homeostasis | LX1.0 | 1 | 1 |
| YJR051W | OSM1 | Protein folding, nucleobase-containing small molecule metabolic process | LX1.0 | 1 | 1 |
| YER115C | SPR6 | Meiotic cell cycle, sporulation | LX1.0 | 1 | 1 |
| YPR107C | YTH1 | mRNA processing | LX0.1 | 1 | 1 |
| YIR036C | IRC24 | No annotated term | LX0.1 | 5 | 5 |
| YDR033W | MRH1 | No annotated term | LX0.1 | 2 | 2 |
| YCL056C | PEX34 | Peroxisome organization | YX1.0 | 1 | 1 |
| YOL150C | N/A | No annotated term | YX1.0 | 1 | 1 |
| YNL165W | N/A | No annotated term | YX1.0 | 1 | 1 |

| | | | | | |
|---|---|---|---|---|---|
| YBR244W | GPX2 | Response to chemical, response to oxidative stress | YX1.0 | 1 | 1 |
| YBL083C | N/A | No annotated term | YX1.0 | 1 | 1 |
| YKL034W | TUL1 | Protein modification by small protein conjugation or removal, proteolysis involved in cellular protein catabolic process | YX1.0 | 1 | 1 |
| YDR435C | PPM1 | Protein alkylation | YX1.0 | 1 | 1 |
| YJL141C | YAK1 | Transcription by RNA polymerase II, response to starvation, response to heat | YX1.0 | 1 | 1 |
| YPR095C | SYT1 | Exocytosis | YX1.0 | 1 | 1 |
| YNL092W | N/A | Protein alkylation | YA0.1 | 1 | 1 |
| YCL040W | GLK1 | Monocarboxylic acid metabolic process, transmembrane transport, nucleobase-containing small molecule metabolic process, carbohydrate metabolic process, generation of precursor metabolites and energy, carbohydrate transport | YA0.1 | 1 | 1 |
| YHR114W | BZZ1 | Cytoskeleton organization, endocytosis, response to osmotic stress | YA0.1 | 2 | 2 |
| YDL236W | PHO13 | Carbohydrate metabolic process | YA0.1 | 1 | 1 |
| YDR205W | MSC2 | Ion transport, cellular ion homeostasis, transmembrane transport | YA0.1 | 1 | 1 |
| YFL033C | RIM15 | Transcription by RNA polymerase II, meiotic cell cycle, regulation of cell cycle, mitotic cell cycle, response to starvation, response to heat | YA0.1, YA1.0 | 1 + 1 | 2 |
| YGR077C | PEX8 | Transmembrane transport, peroxisome organization, protein targeting | YA1.0 | 1 | 1 |
| YNL202W | SPS19 | Meiotic cell cycle, sporulation, monocarboxylic acid metabolic process, lipid metabolic process | YA1.0 | 1 | 1 |
| YGR040W | KSS1 | Response to chemical, regulation of cell cycle, invasive growth in response to glucose limitation, protein phosphorylation, cytokinesis | YA1.0 | 1 | 1 |
| YDR213W | UPC2 | Response to chemical, transcription by RNA polymerase II | YA1.0 | 1 | 1 |
| YNL009W | IDP3 | Monocarboxylic acid metabolic process, lipid metabolic process | YA1.0 | 1 | 1 |
| YJL212C | OPT1 | No annotated term | YA1.0 | 1 | 1 |
| YFR039C | OSW7 | Meiotic cell cycle, sporulation, cell wall organization or biogenesis | YA1.0 | 1 | 1 |
| 55 | | | | | 104 |

**Table S6**. Genotypes for the three parent strains utilized in this study.

| Strain | Genotype |
|---|---|
| BY4741 | MATa his3Δ0 leuΔ0 met15Δ0 ura3Δ0 |
| BY4742 | MATα his3Δ1 leu2Δ0 lys2Δ0 ura3Δ0 |
| BY4743 | MATa/α his3Δ1 leu2Δ0 lys2Δ0 ura3Δ0 |

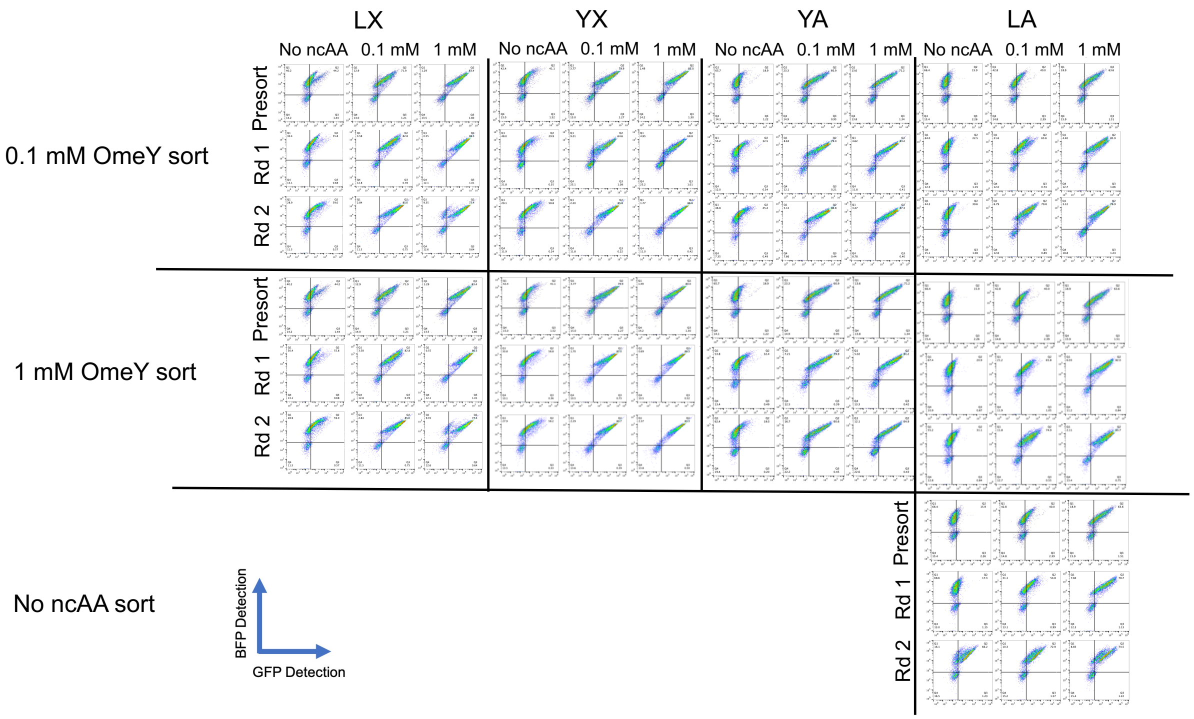


**Figure S1.** Comparison of sorting rounds with analytical flow cytometry.

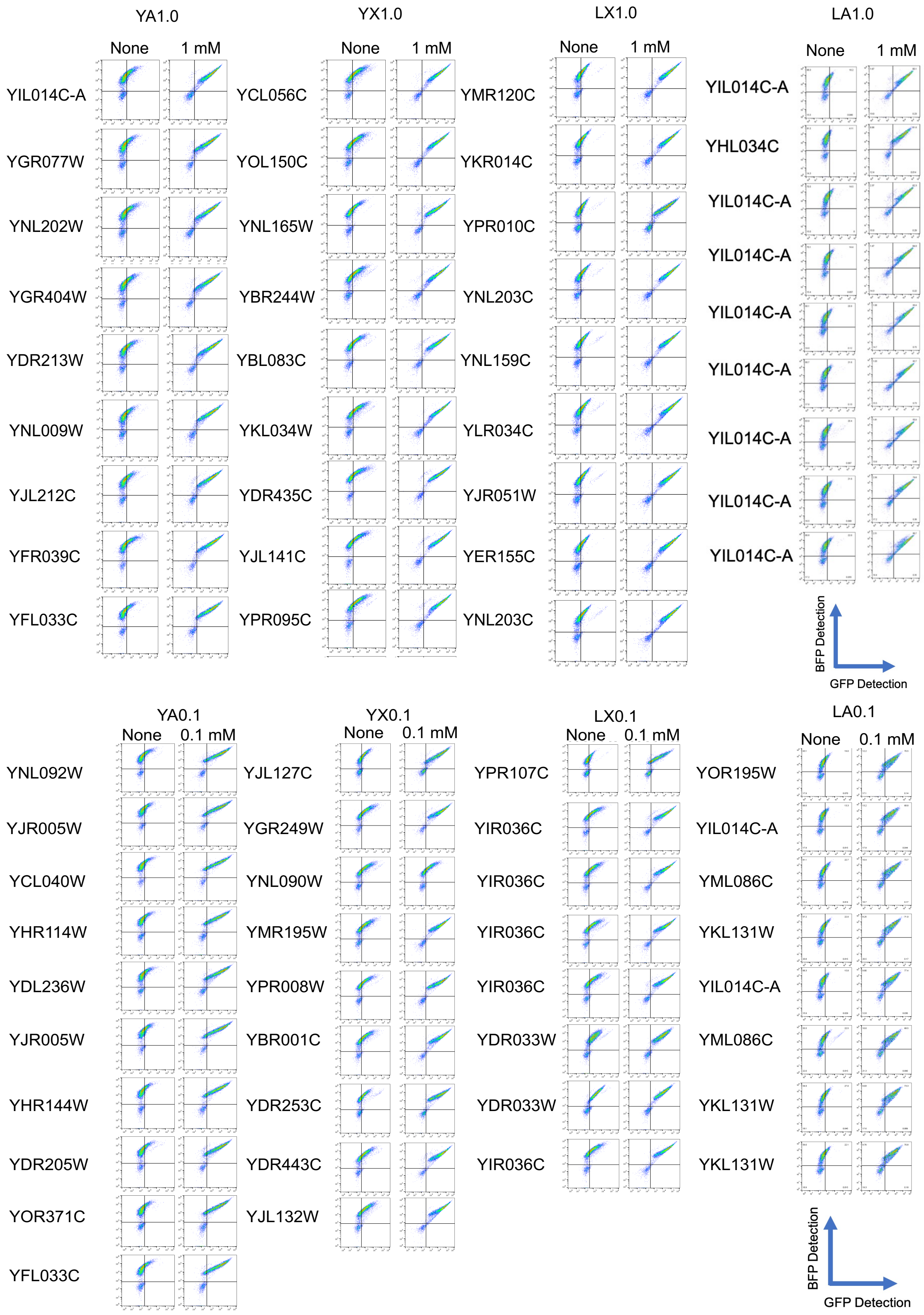
YA1.0
None
1 mM
YIL014C-A
YGR077W
YNL202W
YGR404W
YDR213W
YNL009W
YJL212C
YFR039C
YFL033C
YX1.0
None
1 mM
YCL056C
YOL150C
YNL165W
YBR244W
YBL083C
YKL034W
YDR435C
YJL141C
YPR095C
LX1.0
None
1 mM
YMR120C
YKR014C
YPR010C
YNL203C
YNL159C
YLR034C
YJR051W
YER155C
YNL203C
LA1.0
None
1 mM
YIL014C-A
YHL034C
YIL014C-A
YIL014C-A
YIL014C-A
YIL014C-A
YIL014C-A
YIL014C-A
YIL014C-A
BFP Detection
GFP Detection
YA0.1
None
0.1 mM
YNL092W
YJR005W
YCL040W
YHR114W
YDL236W
YJR005W
YHR144W
YDR205W
YOR371C
YFL033C
YX0.1
None
0.1 mM
YJL127C
YGR249W
YNL090W
YMR195W
YPR008W
YBR001C
YDR253C
YDR443C
YJL132W
LX0.1
None
0.1 mM
YPR107C
YIR036C
YIR036C
YIR036C
YIR036C
YDR033W
YDR033W
YIR036C
LA0.1
None
0.1 mM
YOR195W
YIL014C-A
YML086C
YKL131W
YIL014C-A
YML086C
YKL131W
YKL131W
BFP Detection
GFP Detection

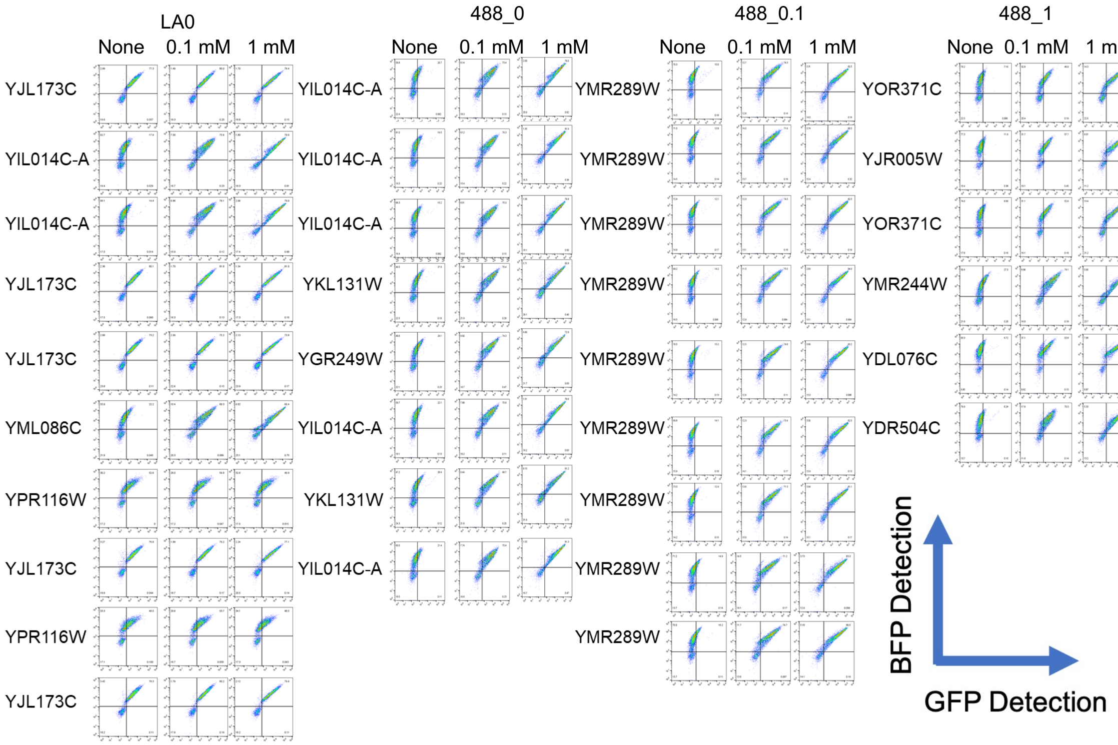


**Figure S2.** Flow cytometry characterization of knockout clones identified following FACS enrichments. All samples were induced in the absence of ncAA and in the same concentration of OmeY used in the sorting path it was identified. Clones identified in the LA0, 488_0, 488_0.1, and 488_1 sorting paths were induced at both 0.1 mM and 1 mM OmeY.

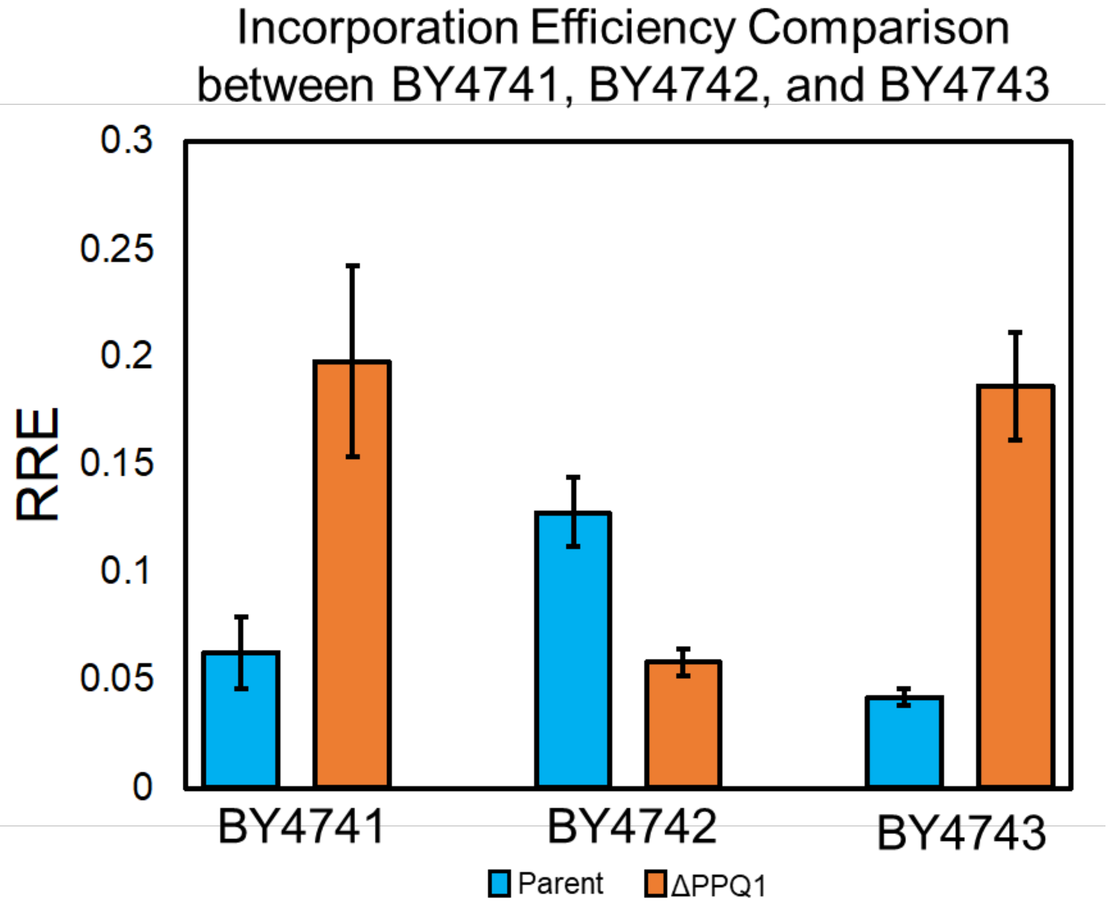


**Figure S3**. Quantitative evaluation of ncAA incorporation efficiency of BY4741, BY4742, and BY4743 parent strains along with *ppq1Δ* knockout strains of each parent. Samples were evaluated in biological triplicate and induced in the presence of 1 mM OmeY. Error bars represent the standard deviation derived from the propagated error.

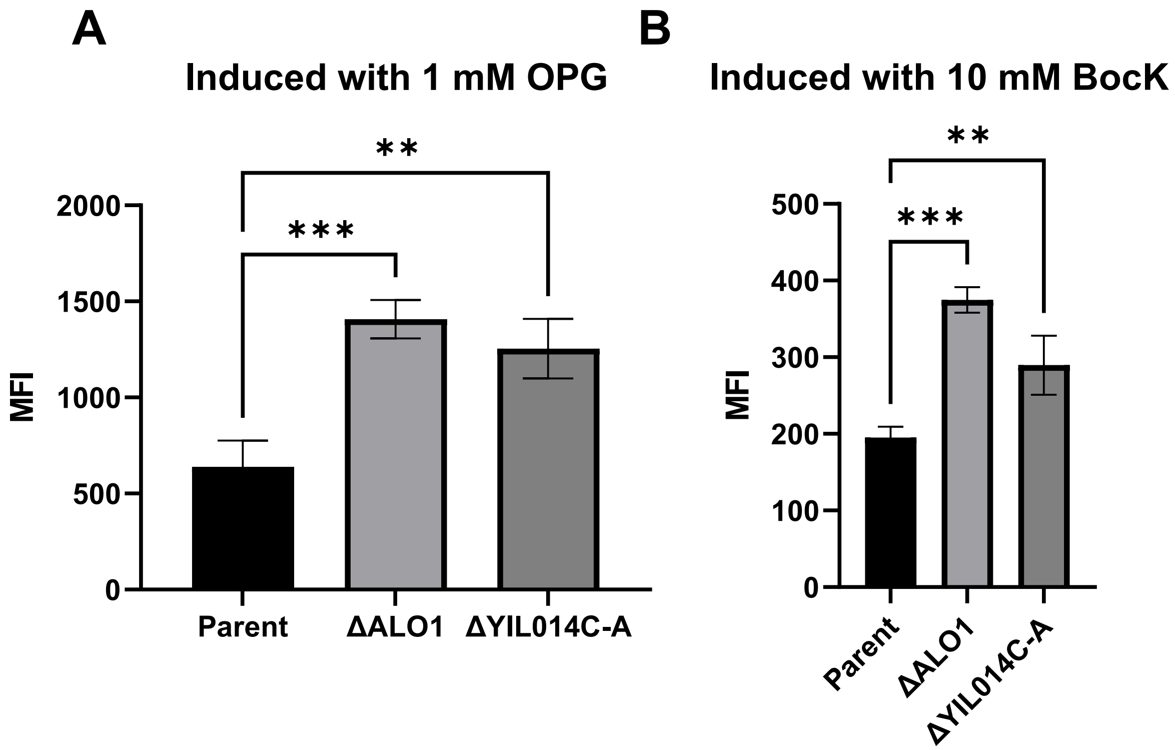


**Figure S4.** One-way ANOVA statistical test using the median fluorescence intensity (MFI) of the C-terminal reporter (GFP) comparing the parent nonmutant strain with knockout strains *yil014c-aΔ* and *alo1Δ* for experiments involving the (A) SpecOPGRS-3 and (B) MaPylRS OTSs. Two stars indicates a *p*-value < 0.01 and three stars indicates a *p*-value < 0.001.

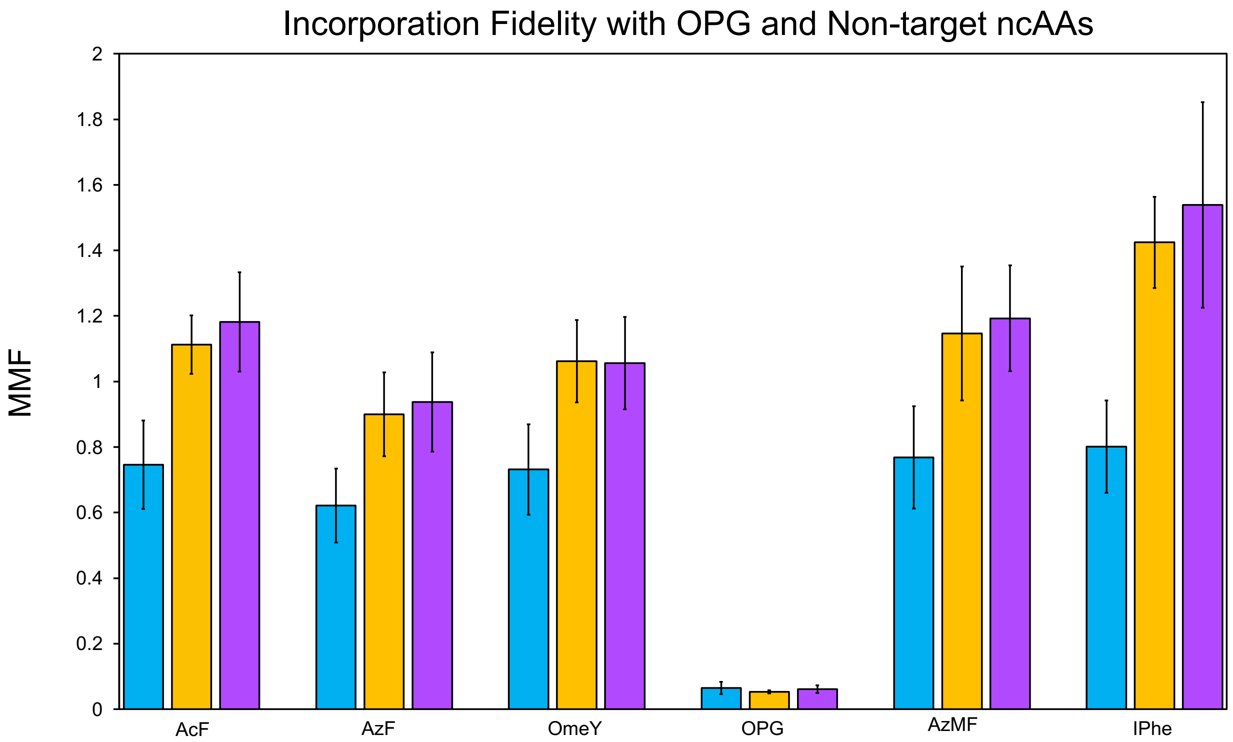


**Figure S5.** MMF values for SpecOPGRS-3 synthetase with OPG and five non-target ncAAs. MMF values greater than or equal to 1 indicate a high frequency of canonical amino acid incorporation.

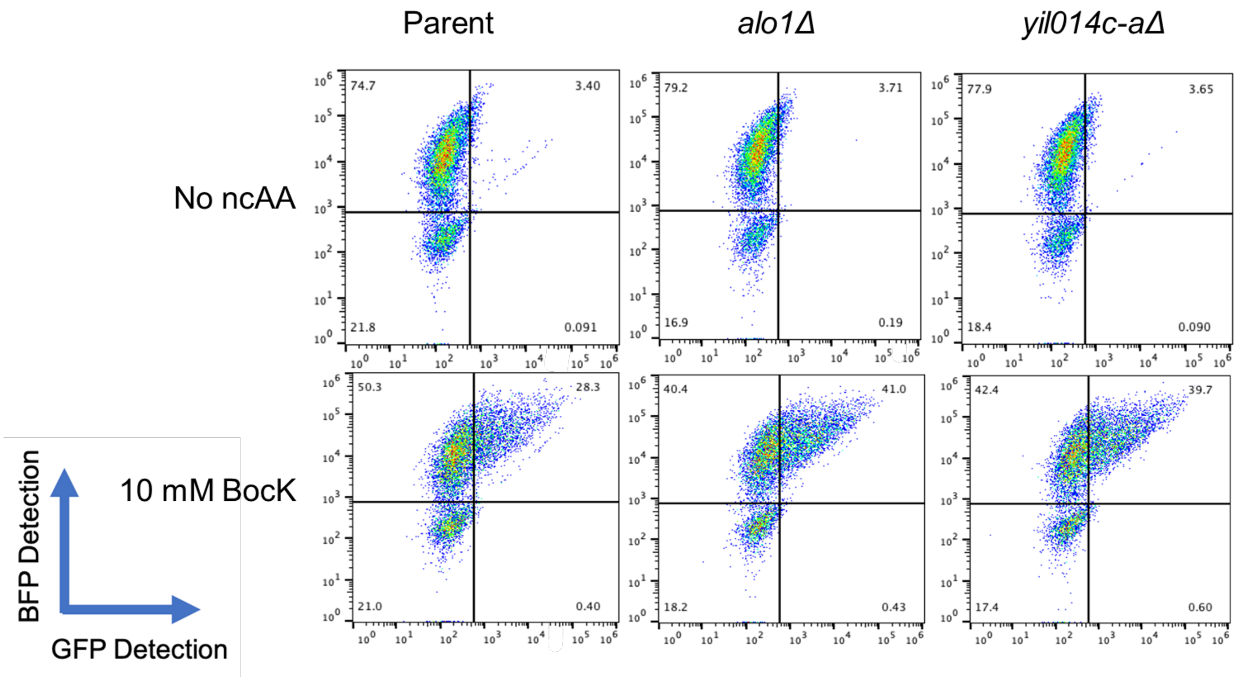


**Figure S6.** Flow cytometry dot plots for MaPylRS experiment. One biological replicate shown from three total replicates for each strain.

$$RRE = \frac{C\ terminus\ detection,\ TAG\ reporter}{N\ terminus\ detection, TAG\ reporter} \Bigg/ \frac{C\ terminus\ detection,\ WT\ reporter}{N\ terminus\ detection,\ WT\ reporter}$$

**SI Equation 1**. Equation for relative readthrough efficiency (RRE).

$$MMF = {}^{RRE_{no\ ncAA}}\!/_{RRE_{ncAA}}$$

**SI Equation 2.** Equation for maximum misincorporation frequency (MMF).
